# ADAR1 p150 prevents HSV-1 from triggering PKR/eIF2α-mediated translational arrest and is required for efficient viral replication

**DOI:** 10.1101/2024.07.29.605561

**Authors:** Adwait Parchure, Mia Cesarec, Vlatka Ivanišević, Marina Čunko, Slađana Bursać, Richard de Reuver, Umberto Rosani, Siniša Volarević, Jonathan Maelfait, Igor Jurak

**Affiliations:** Faculty of Biotechnology and Drug Development, University of Rijeka, Rijeka, Croatia; Department of Molecular Medicine and Biotechnology, Faculty of Medicine in Rijeka, University of Rijeka, Rijeka, Croatia; VIB-UGent Center for Inflammation Research, Ghent, Belgium; Department of Biomedical Molecular Biology, Ghent University, Ghent, Belgium; Department of Biology, University of Padova, Padova, Italy

**Keywords:** ADAR1, HSV-1, protein kinase R PKR, RNA editing, herpesvirus, translational shutdown

## Abstract

Adenosine deaminase acting on dsRNA 1 (ADAR1) catalyzes the deamination of adenosines to inosines in double-stranded RNAs (dsRNA) and regulates innate immunity by preventing the hyperactivation of cytosolic dsRNA sensors such as MDA5, PKR or ZBP1. ADAR1 has been shown to exert pro- and antiviral, editing-dependent and editing-independent functions in viral infections, but little is known about its function in herpesvirus replication. We now demonstrate that herpes simplex virus 1 (HSV-1) hyperactivates PKR in the absence of ADAR1, resulting in eIF2α mediated translational arrest and reduced viral replication. Silencing of PKR or inhibition of downstream signaling pathways by viral (ICP34.5) or pharmacological (ISRIB) inhibitors rescues viral replication in ADAR1-deficient cells. Upon infection, ADAR1 p150 directly interacts with PKR and prevents its overactivation. Our findings demonstrate that ADAR1 is an important proviral factor that raises the activation threshold for sensors of innate immunity.

## Introduction

The adenosine deaminase acting on double-stranded RNA (dsRNA) (ADAR) family of proteins catalyze the conversion of adenosines to inosines in double stranded RNAs (dsRNA) (called A-to-I editing) [1, 2]. A-to-I editing is the most common posttranscriptional RNA modification in mammals which prevents aberrant immune responses to cellular dsRNAs [3–5]. There are three mammalian ADAR proteins: catalytically active ADAR1 and ADAR2, and the catalytically inactive ADAR3 [1]. ADAR1 is predominately expressed as a constitutive, mainly nuclear 110 kDa isoform (referred as p110) and as an interferon (IFN)-inducible 150 kDa isoform (p150) that localizes to both nucleus and cytoplasm [6–8]. ADAR1 is essential for mouse development and regulation of cellular homeostasis [9–11], and in humans, mutations and dysregulation of ADAR1 activity is associated with a number of diseases, including cancer and Aicardi-Goutières syndrome, a severe autoimmune disease associated with spontaneous IFN production (i.e. interferonopathy)[12–14]. The ADAR1 editing function is in conjunction with its major role in the control of innate immunity, namely the prevention of hyperactivation of various cytosolic cellular dsRNA sensors such as retinoic acid-inducible gene I (RIG-I)-like receptors (RLRs), melanoma differentiation-associated protein 5 (MDA5) and laboratory of genetics and physiology 2 (LGP2), 2′–5′-oligoadenylate synthase (OAS), Z-DNA-binding protein 1(ZBP1) and protein kinase R (PKR) (reviewed in [15] [16–22]. These sensors recognize dsRNAs that were previously attributed exclusively to viral infections, as dsRNA is an integral part of their replication cycle (RNA viruses) or the result of transcription of overlapping antisense genes and internal single stranded RNA (ssRNA) secondary structures (RNA and DNA viruses). However, increasing evidence suggests that dsRNA may have an endogenous origin in the transcription of pseudogenes, retroelements, mitochondrial DNA, and transcripts containing Alu elements (a class of repetitive short interspersed elements, IR*Alus*) [23, 24]. The activated dsRNA sensors trigger signals that lead to a less favorable state for viral replication. For example, upon dsRNA binding RLRs interact with mitochondrial antiviral-signaling protein (MAVS) that activates TANK-binding kinase 1 (TBK1) and IκB kinase-ε (IKKε), which in turn activate IRF3 and IRF7; these, together with nuclear factor-κB (NF-κB), then induce type I IFN and other antiviral or immunoregulatory genes [25]. On the other hand, dsRNA-activated OAS impedes translation by synthesizing 2′–5′-linked oligoadenylates and activating the latent endoribonuclease RNase L, which degrades RNA of both cellular and viral origin, leading to inhibition of viral replication [26]. ZBP1 and ADAR1 are the only two mammalian proteins that contain a Z-DNA Zα-domain, and mutation or absence of ADAR1 has been shown to lead to accumulation of Z-RNA, which subsequently activates ZBP1 and induces NF-κB activation (at least in human cells), necroptosis or apoptosis via the receptor-interacting serine/threonine protein kinase – 3 (RIPK3) pathway [27–29] [30]. Finally, dsRNA-dependent protein kinase (PKR) is a major mediator of the antiviral and inflammatory response, and a number of viruses have been found to encode proteins that inhibit PKR signaling. Once activated, PKR phosphorylates the eukaryotic initiation factor eIF2a, which binds the initiator methionyl-tRNA, and sequesters eIF2B, which is critical for recycling eIF2 between successive rounds of initiation, thereby blocking translation and impairing viral replication [31].

The role of ADAR1 has been intensively studied in RNA viruses because of the inevitable dsRNA stage of their replication, and it has been shown that ADAR1 exerts both pro- and antiviral activity, in an editing-dependent and editing-independent manner [32, 33]. For example, ADAR1 plays an important role in suppressing apoptosis, PKR activation and RLR signaling in cells infected with various RNA viruses, including measles virus (MV), influenza A virus (IAV), vesicular stomatitis virus (VSV), flaviviruses, etc. [32, 34–37]. In contrast, the role of ADAR1 in herpesviruses (order *Herpesvirales*), large dsDNA viruses with a complex two-phase replication cycle (i.e. productive and latent phase), is largely unknown and has only been studied in a few cases. Nevertheless, differential expression of ADAR1 in infected cells has been observed in a range of different herpesviruses, from mollusk to human herpesviruses [33]. For instance, the levels of both ADAR1 forms, p110 and p150, increase during KSHV reactivation [38], whereas in cells productively infected with HCMV, only the ADAR p110 form is upregulated [39]. These results indicate that viruses from the same family, impact ADAR1 functions in different ways. Indeed, herpesvirus transcriptome studies, although limited in number and comprehensiveness, show hyper-editing of transcripts that are not conserved, neither in sequence nor function, between different herpesviruses [33]. For instance, selective editing was found for non-conserved miRNAs encoded by KSHV [40, 41], EBV [42, 43] and HSV-1 [44, 45], while on the other hand, editing of host miR-376a was found to be important for HCMV infection [39]. Furthermore, it was recently shown that depletion of ADAR1 in latently infected KSHV cells inhibits viral gene transcription and viral replication during reactivation while increasing IFN induction. Efficient reactivation of KSHV in ADAR1-depleted cells can be rescued by depletion of RIG-I, MDA5 or MAVS, suggesting that ADAR1 serves as a proviral factor by attenuating the response initiated by host pattern recognition receptors (PRRs) [38].

The role of ADAR proteins in HSV-1 infection has not yet been investigated, but HSV-1 miRNAs are known to be edited in latently infected human ganglia [44]. During latency, the viral genome is largely repressed, and only transcripts derived from the latency-associated transcript (LAT) locus are abundantly expressed [46]. The LAT transcripts give rise to a very stable intron and a series of viral miRNAs [47–49], one of which, miR-H2, is hyper-edited [44]. miR-H2 targets a transcript encoding ICP0, a potent transactivator of gene expression and a ubiquitin ligase and which is encoded antisense to the miR-H2 locus [49–51]. Editing of miR-H2 occurs in the seed region of this miRNA and leads to an additional number of potential targets, including ICP4, the major HSV-1 transcriptional regulatory protein [44]. The exact biological significance of this hyper-editing is not yet clear, but it suggests that ADAR proteins are involved in regulation of HSV-1 latency. In contrast to latency, during which transcription is restricted, HSV-1 initiates strong transcription of all its genes during productive infection, which in turn are recognized by various cellular immune sensors and can trigger an antiviral response. However, to preclude their activation and signaling HSV-1 encodes numerous immunomodulatory proteins, including the four proteins, US11, ICP34.5, ICP6 and vhs, which are involved in the direct or indirect suppression of antiviral dsRNA sensor signaling [52, 53]. ICP6, the ribonucleotide reductase large subunit, suppresses caspase 8 mediated apoptosis and prevents activation of ZBP1/RIPK3/MLKL-dependent necroptosis [54–56]. US11 binds directly to dsRNAs and blocks the activation of PKR, RIG-I and OAS [57–60]. The main HSV-1 neurovirulence factor, ICP34.5, has many interactors and ascribed functions, including inhibition of autophagy by interacting with Beclin1, suppression of DNA sensing by interacting with cGAS and STING, inhibition of TBK1, and prevention of downstream PKR signaling by recruiting protein phosphatase 1α (PP1) to dephosphorylate eIF2α and prevent translational arrest [61]. In addition, vhs is an endoribonuclease that triggers the selective degradation of host mRNAs to evade the host antiviral response and facilitate viral infection [62, 63]. Deployment of tegument vhs limits the accumulation of dsRNA in infected cells, and vhs mutants show enhanced activation of PKR [64]. [54–56]

In this study, we investigated the role of ADAR1 protein in productive HSV-1 infection and found that in the absence of ADAR1 viral protein expression and viral replication are reduced. The absence of ADAR1 leads to overactivation of PKR and its downstream signaling, resulting in impaired viral replication. Our results suggest that ADAR1 has an important proviral function by attenuating the activation of dsRNA sensors, highlighting the dsRNA-sensor interfering-function of ADAR1 as a potential target for the development of new antiviral drugs.

## Results

### ADAR1 is required for efficient HSV-1 replication

To investigate the role of ADAR1 in HSV1 infection, we used HEK293 cells deficient for both the p110 and p150 isoforms of ADAR1 (ADAR1 KO) [17, 65]. The ADAR1 KO cells and the parental cells (ADAR1 WT) exhibited normal morphology, comparable duplication time and levels of apoptosis. It is important to mention that we occasionally observed ADAR1-positive cells in the KO cell cultures, as indicated by the appearance of a weak signal for ADAR1 p110 protein in the Western blot (Figure 1C) or some ADAR1-positive cells in immunofluorescence (not shown), which are likely remnants of inefficient ADAR1 CrispR/Cas9 deletion. However, since ADAR1-negative cells constituted more than 99% of the cells in culture at all time points of the experiments, we believe that this did not significantly impact the results of our study.

**Figure 1.**
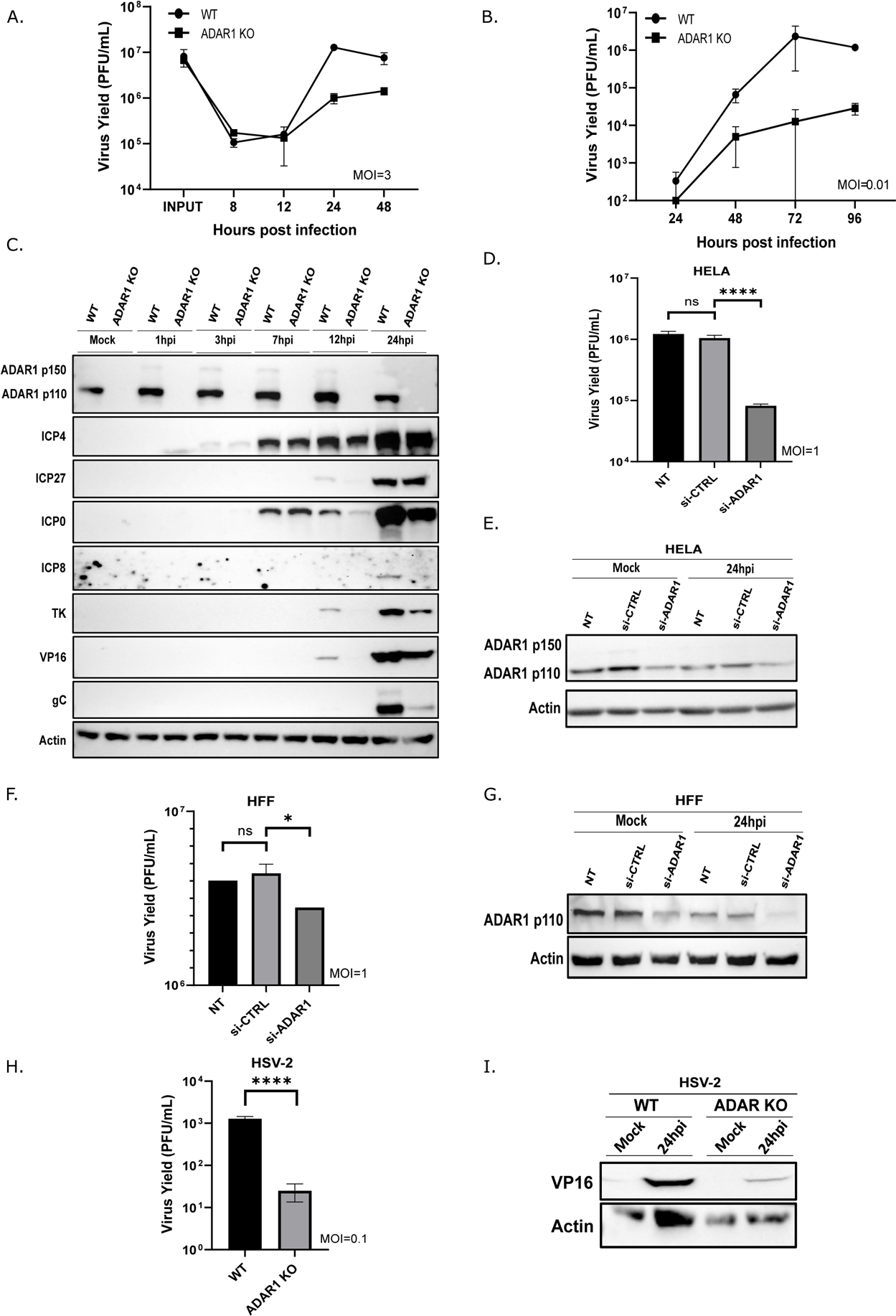
ADAR1 is required for efficient HSV-1 replication. **(A-C)** WT and ADAR1 KO were infected with HSV-1 at MOI of 3 (A) or 0,01 (B) and samples for titration or Western blot analysis (C) were collected at indicated times after infection. **(D-E)** Hela were transfected with control or ADAR1 siRNA for 24 h and then infected with HSV-1 (MOI=1). Virus yield was determined (D) and protein analysed by Western blot (E) 24 h.p.i. **(F-G)** HFF-1 cells were transfected with siRNA and infected with HSV-1 (MOI=1). Virus yield was determined (F) and protein analysed by Western blot (G) 24 h.p.i. **(H-I)** WT and ADAR1 KO cells were infected with HSV-2 (MOI=0.1) and samples for titration (H) or Western blot analysis (I) were collected 24 h.p.i. The data shown is representative of multiple (>3) independent experiments performed in triplicates (A, B, D, F and H) are shown as mean ± standard deviation (SD). ns – no statistical significance; *, p≤0.05; ****, p<0.0001 by One way ANNOVA for (D and F) and Student’s T test for (H).

We first infected ADAR1 WT and ADAR1 deficient cells with HSV-1 strain KOS at high (MOI of 3) and low MOI (0.01) and analyzed viral protein expression and virus yield during the course of infection. Our results show that HSV-1 replication was 10x and > 1000x more productive in ADAR WT than in ADAR1 KO cells at high and low MOI infection, respectively (Fig. 1A and B). Western blot analysis shows that ADAR1 protein levels, both p110 and IFN-inducible p150 isoforms, do not change significantly during the course of infection at high MOI, consistent with the depletion of ADAR1 mRNA (Suppl. Figure 1). This is also consistent with the stable ADAR1 expression levels in cells infected with HSV-1 or HSV-2 observed by Nachmani et al. [39], but in contrast to cells infected with HCMV [39] or with other herpesviruses in which different isoforms of ADAR1 are induced [33]. Furthermore, in most other cells tested, we did not observe an increase but rather a decrease of ADAR1 after infection with HSV-1. (e.g. HeLa, HFF, A549) (Figure 1D-G). Not surprisingly, we could not reproducibly detect ADAR2 protein in ADAR1 WT or KO cells or other cells tested, as ADAR2 expression is mostly restricted to the central nervous system [66]. Our analysis shows that the decreased expression of late viral genes corresponds to the replication defect in ADAR1 KO cells, as indicated by the levels of the late viral protein gC. However, interestingly the initial expression of IE proteins (ICP0, ICP4 and ICP27) was largely comparable in WT and KO cells until 6 hours post infection (h.p.i.) or even throughout the course of infection (i.e. ICP4, Figure 1C). These results indicate that ADAR1 is required for efficient HSV-1 replication and that the block to efficient viral replication occurs at a later stage of infection and not at the entry or initiation of gene expression.

Next, to test whether the presence of ADAR1 is a general requirement for efficient HSV-1 infection and not a specific property of the HEK293 derivative, we knocked down ADAR1 with siRNA in various cell lines and primary cells, including primary HFFs, HeLa, and A549. Indeed, we observed the same reduced HSV-1 replication phenotype in most cell lines, albeit to varying degrees (from 1,5x to > 10x; Figure 1D and F). The observed differences in the requirements for ADAR1 in different cells can be attributed to the degree of ADAR1 downregulation (Figure 1 E and G), but also to biological differences between different cells. In addition, we investigated whether the observed requirement for ADAR1 proteins is only a property of HSV-1 or whether it is also important for the replication of other viruses in the group. To test this, we infected ADAR1 KO cells with herpes simplex virus 2 (HSV-2), a closely related virus that primarily infects the genital area and establishes latency in the dorsal root ganglia, and observed that replication of HSV-2 was also significantly reduced in ADAR1-deficient cells (Figure 1 H and I).

Taken together, our initial results clearly show that the ADAR1 protein is important for the productive replication of HSV-1 and also HSV-2.

### PKR signaling is overactivated in ADAR1 deficient cells

We have shown that ADAR1 plays an important role in HSV-1 infection (Figure 1), and next we wanted to determine the molecular mechanisms of how does ADAR1 exerts its proviral role in HSV-1 infection. First, we wanted to exclude the possibility that HSV-1 infection induces apoptosis in infected ADAR1 KO cells, thereby limiting viral replication. To address this, we infected the cells in the presence of zVAD, a cell-permeant pan caspase Inhibitor of apoptosis, and found that the addition of this pan-caspase inhibitor had no effect on HSV-1 replication in both KO and WT cells (Figure 2A). Nevertheless, we observed a non-significant but slightly increased susceptibility to apoptosis induced by stuarosporin (a microbial alkaloid and potent inhibitor of protein kinases) in ADAR1 KO compared to WT cells, which can be effectively prevented in both cell lines with zVAD (Suppl. Figure 2). Taken together, this led us to conclude that induction of apoptosis is an unlikely limiting factor for HSV-1 replication in ADAR1 KO cells.

**Figure 2.**
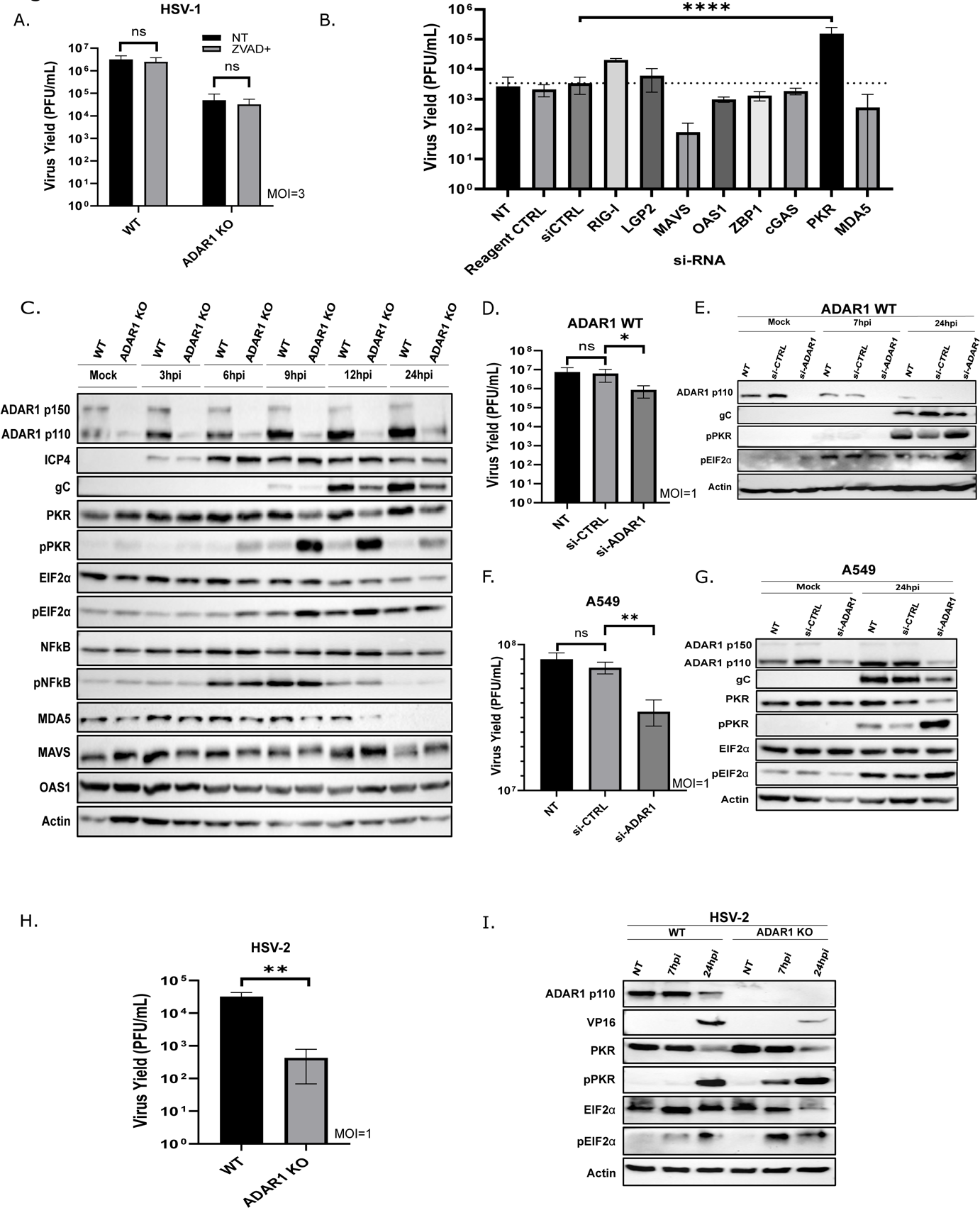
PKR signaling is overactivated in HSV-1 infected ADAR1 deficient cells. **(A)** WT and ADAR1 KO cells were mock treated (NT) or treated with 20μM zVAD for 30min and infected with HSV-1 (MOI=3). One hour after infection, infectious media was removed and fresh media containing zVAD was added. Virus titers in the supernatants were determined 24 h.pi. **(B)** ADAR1 KO cells were transfected with indicated siRNAs and 24 h after infected with HSV-1 (MOI 1). Virus titers in the supernatants were determined 24 h.p.i. Non-treated (NT), transfected transfection reagent only (Reagent CTRL), random scrambled siRNA control (siCTRL). The results represent the mean values of two independent experiments performed in quadruplicates. **(C)** WT and ADAR1 KO were infected with HSV-1 (MOI=3). At indicated times after infection (hpi) cells were collected for western blot analysis. **(D-E)** WT cells were transfected with indicated siRNAs. After 24 hours cells were infected with HSV-1 (MOI=1) and 24 h.p.i. samples for titration (D) and Western blot (E) were collected. **(F-G)** A549 cells were transfected with indicated siRNAs. After 24 hours cells were infected with HSV-1 (MOI=1) and 24 h.p.i. samples for titration (D) and Western blot (E) were collected. Non-treated (NT), scrambled siRNA control (siCTRL), siRNA against ADAR1 (siADAR1). **(H-I)** WT and ADAR1 KO were mock infected (MOCK) or infected with HSV-2 (MOI=1) and samples for the analysis collected 24 h.p.i. The virus yield was determined in the supernatants (H) and protein analyzed by Western blot (I). Data from (A, B, D, F and H) are shown as mean ± standard deviation (SD). *, p≤0.05; **, p<0.01, **** p<0.0001 by One Way ANNOVA for (B, D and F) and Student’s T test for (each pair in A and H).

Next, to reveal the molecular mechanisms behind the viral replication deficit in ADAR1 KO cells we applied siRNA screen targeting well-defined ADAR1 interactors. There is growing evidence that ADAR1 prevents hyperactivation of various cytosolic dsRNA sensors, and thus we reasoned that depletion of effectors responsible for activation of specific host defense mechanisms would lead to complementation of ADAR1 deficiency and rescue HSV-1. Therefore, we transfected ADAR1 KO cells with a series of siRNAs targeting RIG-I, LGP2, OAS1, MDA5, ZBP1, PKR (all dsRNA binding proteins) and MAVS, and assayed virus replication. As a control, we also included a dsDNA binding protein cGAS, which has previously been shown to play an important antiviral role in HSV-1 infection. Initially, we were surprised that the depletion of MDA5 and its downstream effector MAVS had no significant effect on HSV-1 replication. Similarly, depletion of OAS1, ZBP1 and cGAS did not complement ADAR1 deficiency. On the other hand, we observed a slight but reproducible increase in replication when using RIG-I and LGP2 siRNA, (Figure 2B). However, to our surprise we found that only depletion of PKR strongly complemented ADAR1 deficiency and rescued HSV-1 replication (Figure 2B).

To validate our siRNA screen, we compared the expression of a large set of target proteins between ADAR1 WT and KO cells, including the proteins targeted in the siRNA screen, by Western blot during the course of infection (Figure 2C). And indeed, we observed a dramatic difference in PKR activation shown by its autophosphorylation at Thr 446 between KO and parental cells in the early to late phase of infection (6-24 h.p.i.), which coincided with the increased phosphorylation of eIF2α (Ser 51), one of its downstream effectors (Figure 2C), but not with the other, i.e. NF-κB p65 phosphorylation (Ser 536). These results complement the results of the siRNA screen and together with the exclusion of apoptosis, suggest that the mechanism responsible for the less efficient replication of HSV-1 in ADAR1-deficient cells is PKR-mediated repression of general protein synthesis. Important to note, the expression of MAVS, MDA5, and OAS, the other proteins included in the screen, was comparable between the two cell lines, but we did not detect expression of RIG-I, LGP2, ZBP1, RIPK3 and cGAS, indicating low expression or absence of these proteins in both WT and KO cells (Figure 2). It is possible that such a proteome composition permits establishment of ADAR1-KO cells, but the exact contribution of these proteins to the observed phenotype we could not adequately asses. Or in other words, our results suggest that, at least in our model, these proteins are not responsible for the observed phenotype.

A similar association between PKR activation and reduced viral replication in the absence of ADAR has been observed for several RNA viruses, which is consistent with our observation. Nevertheless, we wanted to test whether ADAR1-mediated suppression of PKR activation in productive HSV-1 infection is a general property of infected cells and not a specific property of ADAR1-KO cells derived from ADAR1-WT-HEK293 cells. To address this question, we transfected ADAR1 WT cells as well as human lung carcinoma A549 cells with siRNA targeting ADAR1. Our results show that depletion of ADAR1 in these cells not only reduces viral replication but also triggers enhanced activation of PKR (Figure 2D - G). Furthermore, we show increased activation of PKR signaling in ADAR1-deficient cells infected with HSV-2, suggesting an conserved molecular mechanism at least at the level of the *Simplexvirus* genus (Figure 2 H and I). Our results show that PKR signaling is the primary signaling pathway activated in ADAR1-deficient cells and that downstream signaling may be responsible for limiting viral replication.

### Depletion of PKR or reversal of its negative effects on eIF2**α** rescues HSV-1 replication in ADAR1 deficient cells

Our siRNA screening and Western blot suggest activation of PKR and PKR/eIF2α-mediated suppression of HSV-1 replication in ADAR1-depleted cells. To establish a direct link between PKR and the phenotype, we performed an additional set of siRNA experiments. First, we depleted PKR in ADAR1 WT and ADAR1 KO cells using siRNA and analyzed signaling and viral yield. In both cells, we were able to successfully deplete PKR protein (Figure 3A), which is also consistent with the reduced levels of phosphorylated form of PKR upon infection. This was particularly evident in PKR-depleted KO cells compared to control siRNA. Reduced PKR phosphorylation was accompanied by increased expression of the late protein gC (Figure 3A) and increased viral replication (Figure 3B), which was comparable to ADAR1 WT cells treated with control siRNA. We also observed slightly increased viral replication in PKR-depleted ADAR1 WT cells, suggesting that limited but inhibitory PKR activation also occurs in the presence of ADAR1 (Figure 3B), consistent with observations by other researchers [67]. Next, to show that rescue of HSV-1 replication in the absence of ADAR1 by depletion of PKR is a general property, and not limited to ADAR1 KO cells and its specific proteome, we simultaneously depleted ADAR1 and PKR in unrelated A549 cells (Figure 3C). We observed, as expected, that depletion of ADAR1 leads to activation of PKR and reduced replication (see Figure 2G and Figure 3C). On the other hand, infection of cells with simultaneous depletion of both proteins results in replication comparable to that of untreated cells (Figure 3D). Furthermore, simultaneous siRNA targeting of RIG-I, LGP2, MAVS, OAS, ZBP1, MDA5 or cGAs together with depletion of ADAR1 failed to rescue viral replication in A549 cells. These results clearly indicate a general importance of the ADAR1/PKR interaction for productive HSV-1 infection, but the exact mechanism and downstream effectors were not clear.

**Figure 3.**
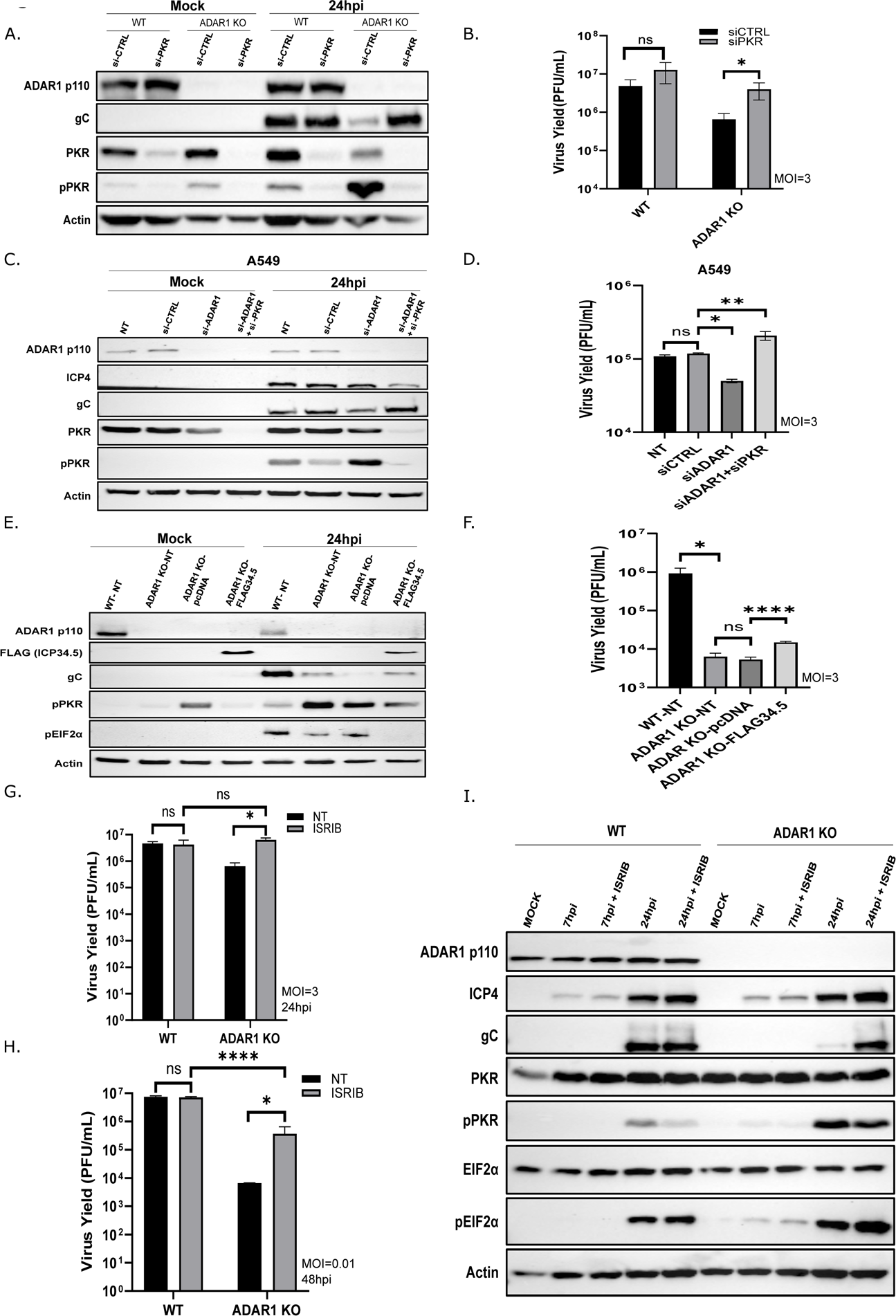
Depletion of PKR or inhibition of its downstream signaling rescues HSV-1 replication in ADAR1 KO cells. **(A-B)** WT and ADAR1 KO were mock transfected or transfected with the indicated siRNA and 24 hours later infected with HSV-1 (MOI=3). (A) Western blot analysis of proteins extracted from infected cells 24 h.p.i. (B) Virus titer in supernatants collected at 24 h.p.i. **(C-D)** A549 cells were transfected with the indicated siRNA and 24 hours later infected with HSV-1 (MOI=3). (C) Western blot analysis of proteins extracted form infected cells 24 h.p.i. (D) Virus titer in supernatants collected at 24 h.p.i.. **(E-F)** ADAR1 KO cells were mock transfected or transfected with a control plasmid (pcDNA) or plasmid expressing FLAG-ICP34.5 protein. 24 h.p. transfection cells were infected with HSV-1 (MOI=3). (E) Western blot analysis of proteins extracted from infected cells 24 h.p.i. (F) Virus titer in supernatants collected at 24 h.p.i. **(G-I)** WT and ADAR1 KO cells were pre-treated with 0.5μM Integrated Stress Response Inhibitor (ISRIB) for 30min, and infected with HSV-1 (MOI=3 or 0.01). (G) Cells infected with HSV-1 (MOI=3) were collected for western blot at 24 hpi. (H) Virus titer in supernatants collected from cells infected with HSV-1 at an MOI of 3 at 24hpi. (I) Virus titer in supernatants collected from infected cells with HSV-1 at an MOI of 0.01. (B, D, F, H, I) The data shown is representative of multiple (>3) independent experiments performed in triplicates. Data from (B, D, F, G and H) are shown as mean ± standard deviation (SD). *, p≤0.05; **, p<0.01, **** p<0.0001 by Student’s T test in each comparison for (B, in F WT-NT to KO-NT, G and H) and One Way ANNOVA for (D and F).

Concomitant with the activation of PKR, we observed phosphorylation of eIF2α (Figure 2C), indicating inhibition of general protein synthesis. This result also suggests that inhibition of downstream PKR signaling on eIF2α or evasion of translational block in ADAR1 KO cells would also rescue HSV-1 replication. To test this, we applied genetic and pharmacological approach. In first, we transfected KO cells with a plasmid expressing ICP34.5, the main HSV-1 neurovirulence factor which recruits protein phosphatase 1 (PP1) to dephosphorylate eIF2α and prevents PKR-triggered translational block [68]. The presence of ICP34.5 in KO cells prior to infection successfully reduced the phosphorylation of eIF2a and PKR, resulting in higher expression of the viral protein gC (Figure 3E). Furthermore, replication of HSV-1 was slightly, but significantly more efficient in KO cells expressing ICP34.5 than in cells transfected with a control plasmid or in non-transfected cells (Figure 3F). It is important to note that we observed a slight but reproducible complementation of ADAR1 deficiency by IPC34.5, which we attribute to the low transfection efficiency in these experiments (i.e., we estimate that less than 30% of cells were transfected) and therefore likely represents an underestimation of the true efficiency of ICP34.5 in inhibiting this pathway. To further substantiate the notion that PKR/eIF2a-mediated translational block is limiting viral replication, we treated KO cells with ISRIB, a pharmacological inhibitor of integrated stress response. The addition of ISRIB reverses the effects of eIF2α phosphorylation and releases translation block by preventing inhibition of eIF2B and maintaining translation initiation [69]. Compared to complementation with proteins expressed by transfected plasmids, we were able to treat all cells and observe more substantial effects by using small molecules. We therefore infected ADAR1 KO cells at low and high MOI in the presence of 500 nM ISRIB and measured viral yield. Clearly, addition of ISRIB effectively rescued the ADAR1 deficient phenotype, at both low and high MOIs (Figure 3 G-I). Surprisingly, however, we observed reduced phosphorylation of PKR but not reduced eIF2α phosphorylation, in infected cells treated with ISRIB compared to infected, non-treated or DMSO-treated cells, in both ADAR1 WT and KO cells (Figure 3 G). Thus, although the addition of ISRIB complemented ADAR1 deficiency (Figure 3 G-I) and prevented translational arrest, there may be additional mechanisms by which ISRIB rescues HSV-1 replication in ADAR1 KO cells. Nonetheless, this series of experiments suggests that PKR/eIF2α-mediated translational arrest is the major pathway responsible for reduced viral replication in ADAR1-KO cells.

### Transcription of viral genes is not affected early in infection

In our previous experiments, we established a direct link between PKR/eIF2α-mediated translational arrest and replication defect in infected ADAR1 KO cells, although the exact mechanism that proceed virus replication deficit was not clear. We observed severely restricted expression of late viral proteins and reduced viral replication (Figure 1 and 3) in KO cells. On the other hand, we observed that IE gene products were less affected at least at the beginning of the infection (Figure 1), especially ICP4, which was unaffected throughout the infection and was comparable between WT and KO cells. This may indicate that translation of ICP4 transcripts are not sensitive to eIF2α-mediated repression, similar to activating transcription factor 4 (ATF4), whose translation actually increases under integrated stress response [70, 71]. We therefore wanted to test whether PKR/eIF2α activation actually causes the infected cells to undergo a certain degree of translational arrest, which limits viral protein levels and subsequent replication. We hypothesized that we would observe increased levels of ATF4 in infected ADAR1-KO cells compared to infected parental cells. Indeed, interestingly, we observed slightly increased ATF4 levels in mock-infected ADAR1 KO cells compared to WT cells, which also correlates with somewhat increased phospho-PKR levels in these cells (Figure 2 and 4A). Nevertheless, we observed a slight increase in ATF4 protein levels during infection in both infected ADAR1 WT and infected KO cells compared to mock-infected cells, but in KO cells ATF4 expression increased dramatically towards the end of infection, further indicating translational arrest (Figure 4A). The expression of ATF4 was readily detectible in both cell lines after treatment with thapsigargin, a widely used ER stress inducer [72], which served as a positive control for the stress induction (Figure 4 A). Remarkably, we did not observe an increase in ATF4 transcript abundance, but rather a slight decrease in both cell lines, which is comparable to previous studies (Suppl. Fig. 3).

**Figure 4.**
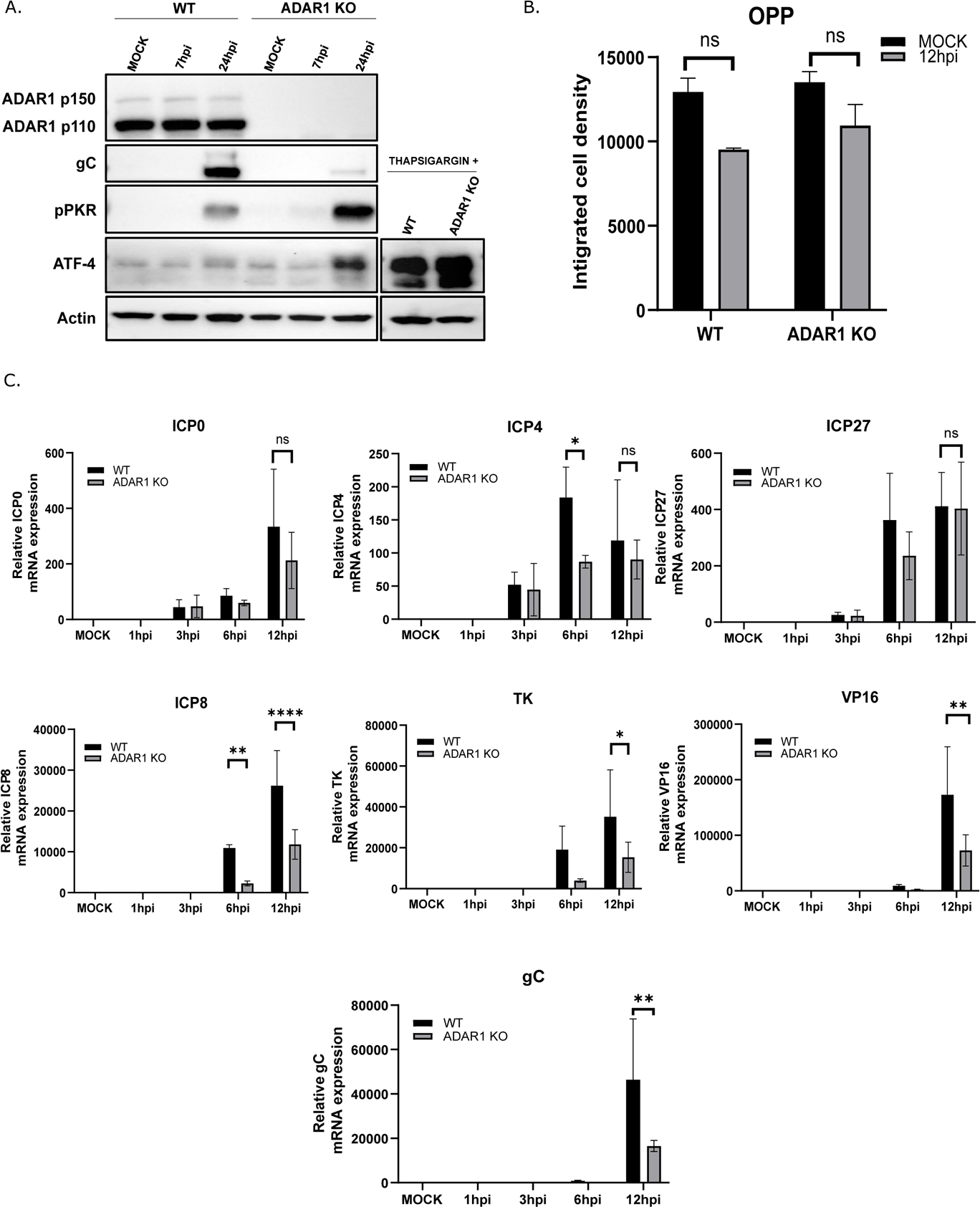
Transcription of viral genes is not affected early in infection. **(A)** ADAR WT and ADAR1 KO were infected with HSV-1 at MOI=1 or treated with thapsigargin for 7 h. At the indicated times p.i. cells were collected western blot analysis. **(B)** WT and ADAR1 KO cells were infected with HSV-1 MOI=1. After 1 hr infectious media was replaced with fresh media. 11 h.p.i. cells were treated with OPP and analyzed using Zeiss LSM700 confocal laser scanning microscope (Carl Zeiss) as previously described [89]. Mean immunofluorescence signal intensity per cell was obtained by dividing the total fluorescence intensity (n = 8–10 fields with 1,500 cells) by the number of cells. **(C)** WT and ADAR1 KO were infected with HSV-1 at MOI=1 in triplicates. RNA was extracted from cells at indicated times after infection (hpi). Immediate early (ICP0, ICP4, and ICP27), early (ICP8 and TK), and late (VP16 and gC) transcripts were detected using RT-qPCR. All samples were normalized to Mock 18S followed by normalization to 1hpi for viral genes. Relative expression of viral genes is indicated in proportion to transcripts detected at 1hpi. Data are shown as mean ± standard deviation (SD), ns not significant, * p≤0.05; **, p<0.01, **** p<0.0001 by Student’s T test for (B) and Two Way ANNOVA for (C).

Next, in an effort to determine the level of translational arrest in infected ADAR1 KO cells compared WT cells, we measured nascent protein synthesis at 12 h.p.i. in infected ADAR1 WT and KO cells using a non-radioactive O-propargyl-puromycin labeling coupled with fluorescent microscopy. As expected, global protein synthesis in mock infected cells was comparable between these two cell lines. On the other hand, total protein synthesis was significantly reduced in both cells, but to our surprise comparable between infected WT and KO cells at 12 h.p.i. Although this result is very intriguing and could support the hypothesis of a potential specific transcripts translational arrest, i.e. affecting only viral or specific transcripts, we are very cautious about drawing such conclusions due to the technical challenges involved. First, the experiments were performed at a relatively high MOI (MOI of 1) to allow for synchronous infection, where differences in viral replication are not substantial and the resolution of the assays may not be sufficient to detect subtle differences.

Our analysis of the viral proteins suggests that their transcription may not be affected, at least in the early phase of infection and as long as these transcripts do not activate the dsRNA sensor PKR and the subsequent translational block. To analyze in detail the events preceding translational arrest, we analyzed the transcription of IE (ICP0, ICP4, ICP27), E (ICP8 and TK) and L transcripts (gC and Vp16) during the course of infection in ADAR1 WT and KO cells at an MOI of 1. Indeed, we observed stalling of all gene classes in the late phase of infection (12 hours), especially of L gene transcripts. In contrast, we observed no statistically significant differences in IE gene transcription until 6 hours post-infection (Figure 4), i.e. around the time of viral DNA synthesis, which also coincides with detectable activation of PKR, suggesting that viral transcripts activate PKR when ADAR1 is absent.

### ADAR1 p150 but not p110 prevents PKR activation in HSV-1 infection

To determine the exact mechanism of proviral ADAR1 activity, it is crucial to identify which of the ADAR1 isoforms, the constitutively expressed nuclear p110 or/and the IFN-inducible cytoplasmic p150 isoform, have the main proviral function and whose deficiency is responsible for the observed phenotype. To address this, we applied two experimental strategies. First, we transfected KO cells with plasmids expressing p110-GFP or p150-GFP proteins and tested the complementation of ADAR1 deficiency for HSV-1 infection. It is important to note that a) up to 30% of cells were successfully transfected (estimate based on GFP expression); b) the levels of ADAR1 proteins expressed from the plasmids were much higher than the endogenous levels of ADAR1 proteins (Figure 5); c) ADAR1 p110-GFP and ADAR1 p150-GFP were predominantly expressed in nucleus and cytoplasm, respectively. Transfected cells were infected with HSV-1 at low (MOI of 0,01) and high MOI (MOI of 3) and analyzed for virus production. Our results show that the ectopic expression of ADAR1 p150, but not p110 or a control plasmid expressing GFP, complements ADAR1 deficiency at both MOIs; virus replication increases for 10x and 2x, compared to control plasmid-transfected cells, at MOI 0.01 and MOI 3, respectively (Figure 5 A-D). Furthermore, these results were supported by analyzing PKR activation upon synchronized infection by Western blot, where we showed that the presence of cytoplasmic ADAR1 p150 isoform, but not nuclear p110, reduced PKR activation (Figure 5 B).

**Figure 5.**
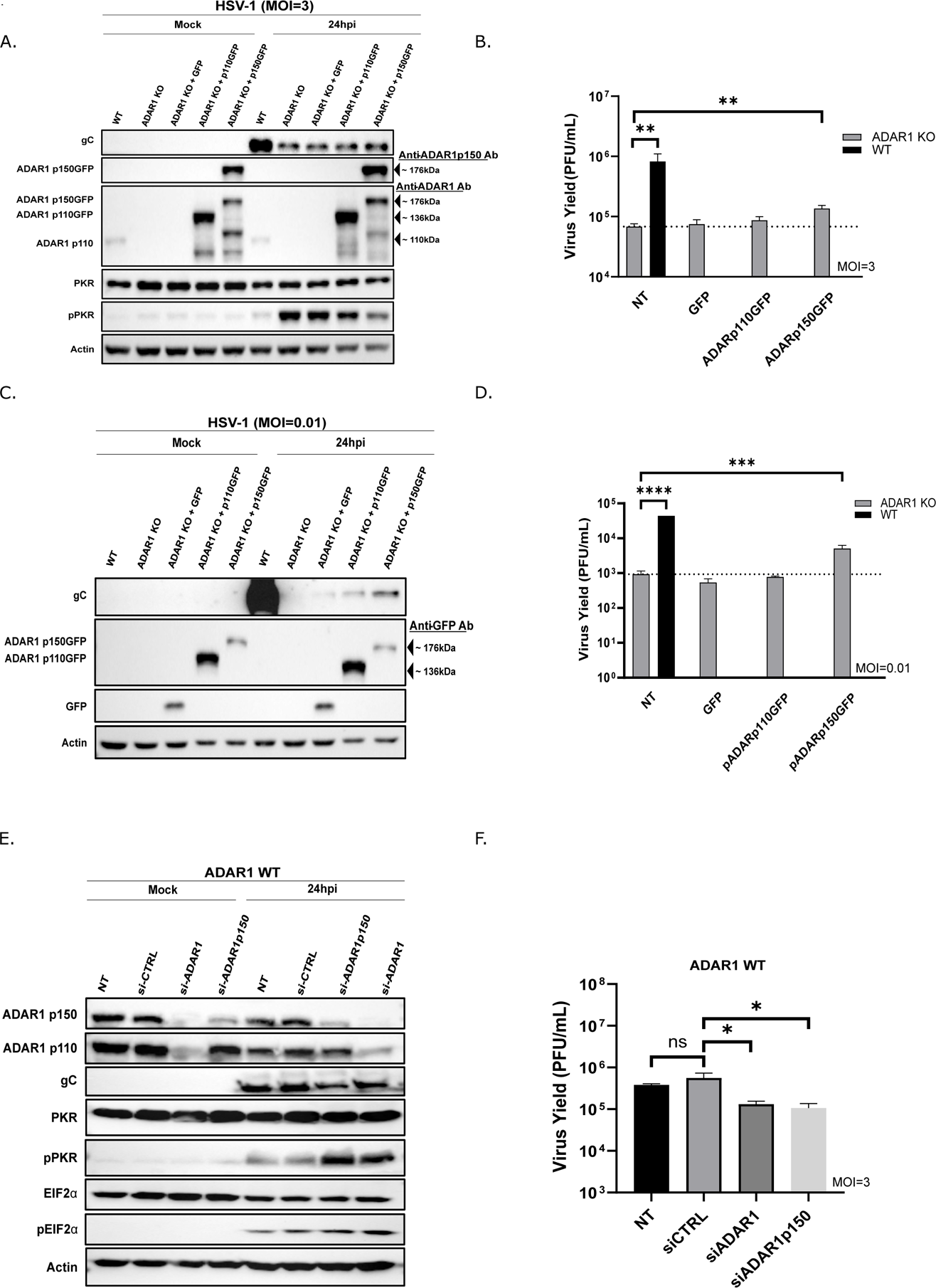
ADAR1p150 and not ADAR1p110 prevents PKR activation in HSV-1 infection. **(A-D)** ADAR1 KO cells were transfected with control plasmid pEGFP-N1 expressing EGFP, and plasmids ADAR1p110GFP or ADAR1p150GFP expressing p110 and p150 forms of ADAR1, respectively, and infected with HSV-1 at the indicated MOI (MOI=3 or MOI=0,01). (A) Cells infected at MOI of 3 were collected at 24 h.pi. and proteins were analyzed by western blot using indicated antibodies to the left of the panels (B) Virus titer in supernatants collected from cells infected with HSV-1 at MOI of 0.1 at 24hpi. (C) Cells infected at MOI 3 were collected at 24 h.pi. and proteins were analyzed by western blot. (D) Virus titer in supernatants collected from cells infected with HSV-1 at MOI 3 at 24hpi. **(E-F)** WT cells were transfected with indicated siRNA and after 24h infected HSV-1 (MOI=3). (E) Cells were collected at 24 h.pi. and proteins analyzed by western blot. (F) Virus titer in supernatants collected at 24hpi. Data are shown as mean ± standard deviation (SD). *, p≤0.05; **, p<0.01, **** p<0.0001 by Student’s T test for independent comparison of WT-NT and ADAR1 KO-NT in (B and D) and One Way ANNOVA for (B, D and F).

To further confirm the role of ADAR1 p150 in productive HSV-1 infection, we used siRNAs to deplete both forms of ADAR1 or selectively the p150 form in ADAR1 WT (Figure 5 E) and A549 (Suppl. Fig 4) cells and tested HSV-1 replication. As expected, depletion of both p110 and p150 and also selective depletion of p150 resulted in increased activation of PKR and consequently decreased HSV replication, confirming that p150 indeed plays a dominant proviral role in HSV-1 infection (Figure 5 E and F).

### HSV-1 infection triggers direct ADAR1 PKR association

We have shown that ADAR1 is important for the prevention of PKR activation and efficient viral replication, but the exact interplay between these two proteins in HSV-1 infection is unclear. Recently, ADAR1 was shown to prevent PKR activation in cells treated with IFN or infected with RNA viruses by direct association with PKR [19, 73, 74]. This led us to hypothesize that HSV-1 or/and host transcripts trigger the association of PKR and ADAR1 at the onset of infection. To address this, we immunoprecipitated PKR in MOCK infected cells, cells treated with IFN and cells infected with HSV-1 at the 7 h.p.i. (i.e. approximately at the time post-infection when we observed the highest PKR activation). PKR co-immunoprecipitated ADAR1 p150 in all tested samples, but the levels of co-immunoprecipitated ADAR1 were significantly higher in HSV-1 infected cells (Figure 6 A and B). Consistent with siRNA p110 specific knockdown and the experiments with ectopic expression of p110, which together suggest a dominant functional role of p150, we were unable to detect the p110 isoform of ADAR1 in immunoprecipitations. Unfortunately, we were not able to perform the reciprocal immunoprecipitation using available ADAR1 p150 antibodies probably due to lower specificity of these antibodies. Nonetheless, our results strongly indicate a direct ADAR1p150-PKR association during HSV-1 infection. Moreover, these results provide further evidence that a deficiency of the p150 form, but not p110, is responsible for the replication deficit in ADAR1 KO cells.

**Figure 6.**
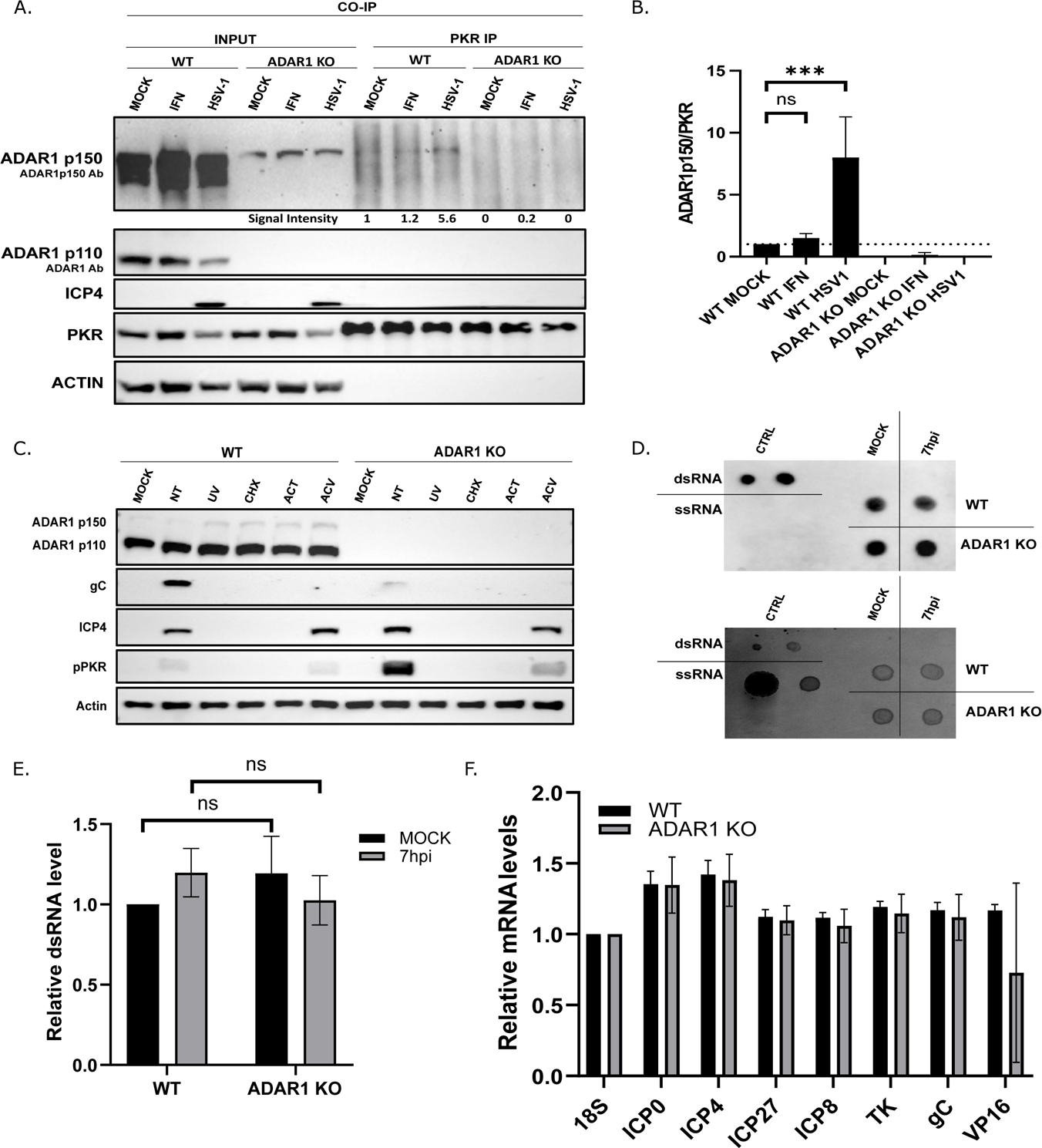
HSV-1 infection triggers direct ADAR1-PKR association. **(A-B)** WT and ADAR1 KO cells were infected with HSV-1 (MOI=3) or treated with 10ng/mL IFN and at 7h.p.i. cells were collected for protein co-immunoprecipitation (CO-IP) using anti-PKR antibody. (A) Representative western blot analysis of the input protein sample and PKR-co-immunoprecipitated proteins. Proteins detected with specific antibodies (indicated in brackets) are shown to the right of the panel. The signal intensities determined with ImageJ are shown below the panel (B) ImageJ quantification of co-immunoprecipitated ADARp150 normalized to the PKR signal. Quantification represents three individual experiments (n=3). **(C)** WT and ADAR1 KO cells were pre-treated with 100ug/mL Cycloheximide (CHX) or 1 g/mL Actinomycin D (ACT) or 100 M Acyclovir (ACV) for 1 h and infected with HSV-1 or UV inactivated HSV-1 at MOI 3. At 24h.p.i. cells were collected for the Western blot analysis. **(D-E)** WT and ADAR1 KO were infected with HSV-1 (MOI=3) and 7h.p.i. RNA was extracted. (D) 1μg of total RNA extracted from infected cells was spotted on a membrane together with 0.01μg and 0.1μg of 142 bp dsRNA (upper left corner of the panel) Jena-Bioscience) and 10μg and 1μg of Poly(A) ssRNA (lower left corner of the panel, Jena-Bioscience). dsRNA was detected using J2 antibody (upper panel). To normalize the loading of the tested samples, the membranes were stained in 0,01% solution of Methylene-blue (lower panel). (E) ImageJ quantification of dot blot normalized to the loading signal. Experiments were performed in multiplicate and shown as mean values (n=18). **(F**) WT and ADAR1 KO cells were infected with HSV-1 (MOI=3) and 7h.p.i. PKR was immunoprecipitated and RNA was extracted from anti-PKR antibody co-immunoprecipitation (CO-IP) samples. Indicated target genes were detected using RT-qPCR and normalized to 18S of corresponding cell type. Experiments was performed in quadruplets. Data is shown as mean ± standard deviation (SD); ns – not statistically significant; ***, p<0.001, by One Way ANNOVA for (B) and Student’s T test for (each pair in E).

### Immediate early or/and early transcripts are required for activation of PKR

Finally, we were very intrigued by the ADAR1-PKR association triggered by HSV-1 infection, and therefore we should endeavor to understand the molecular determinants of this association. It is well established that dsRNAs trigger PKR activation, but the dynamics of these events and whether viral or host transcripts or both trigger this activation are unclear. Our analysis of viral transcripts shows that transcription and translation of viral gene products is comparable between WT and KO cells at the onset of infection, whereas the expression of most viral genes was severely affected by a translational arrest late in infection (Figure 4C). These results suggest that events prior to viral DNA replication cause deleterious activation of PKR and that inhibition of viral DNA replication should not prevent PKR activation. To test this hypothesis and to further characterize functions required for PKR activation, we infected cells treated with acyclovir (ACV; an inhibitor of DNA synthesis and late gene expression), actinomycin D (AcD; an inhibitor of RNA poly II), cycloheximide (CHX; an inhibitor of protein synthesis), or UV inactivated virus. In short, our results show, as expected, that virus attachment or penetration alone is not sufficient to trigger the inhibitory signals (as indicated by UV-inactivated viruses), and that mRNA and protein expression are required for PKR activation (as indicated by treatment with AcD and CHX, Figure 6 C). On the other hand, activation of PKR in ACV-treated cells was clearly detectable in both WT and KO cells, albeit at lower levels than in untreated cells, suggesting that IE and E transcripts or/and proteins are sufficient for initial activation. Progression of infection and replication of its DNA later in infection further increased PKR activation. At this stage, although very intriguing to us, we cannot dissect whether PKR mediated translational inhibition affects different genes or classes of genes differently, or whether a wide-ranging translational inhibition causes a replication defect.

Overall, it is clear that viral transcription is required for PKR activation, but we could not exclude the possibility that induced host transcripts activate or contribute to PKR activation. Recently, it has been demonstrated that host pseudogene transcripts play a role in the activation of the RIG-I signaling pathway in HSV-1 infected cells. To address this question, we first wanted to rule out the possibility that the total amount of dsRNAs that can be targeted by PKR is higher in KO cells due to the lack of ADAR1 editing leading to PKR activation. To investigate this, we measured dsRNA levels with a dot-blot assay (Figure 6D) and with immunofluorescence using the dsRNA binding J2 antibody. Briefly, WT and KO cells were infected at a high MOI and total RNA was extracted 7 h.p.i. (about the time of detected PKR activation) and spotted onto a charged membrane, and dsRNAs were detected with the J2 antibody. Our results suggest that reactive-dsRNA levels in infected KO cells are similar to those in WT cells. This led to the hypothesis that a massive influx of viral transcripts with dsRNA structures leads to activation of PKR and that in the absence of ADAR1 this signaling is exaggerated and limits viral protein synthesis and replication. To confirm that PKR binds viral transcripts, we performed an RNA immunoprecipitation (RIP) assay in which we pulled down PKR and analyzed the RNAs in the complex by RT-qPCR. Our results show that all classes of viral transcripts (i.e. represented with ICP0, ICP4, ICP27, ICP8, TK, gC and VP16) present in infected cells can be found in complex with PKR (Figure 6F), consistent with these transcripts playing a role in PKR activation. Nevertheless, although it is known that in HSV-1 infection the vast majority of host transcripts are depleted by the function of vhs, based on this very limited analysis we cannot exclude the possibility that certain host transcripts also contribute to PKR activation. To fully clarify this question, the associated RNAs need to be sequenced, which is beyond the scope of this manuscript.

Overall, in this study, we comprehensively investigated the proviral function of ADAR1 based on its capacity to limit PKR activation and downstream signaling, which in its absence would lead to translational arrest and inhibition of productive HSV-1 infection. The proviral nature of ADAR1 makes it clear that ADAR1 is a potential target for the development of new drugs against important human infections.

## Discussion

In this study, we show that ADAR1 p150 plays an important role in productive HSV-1 infection, primarily by modulating PKR activation and downstream signaling that prevents translational arrest and enables efficient viral replication. ADAR1 binds dsRNAs and catalyzes the deamination of adenosine to inosine, which is an important marker for the differentiation of self and non-self RNAs and the prevention of an abnormal immune response [75]. Recently, ADAR1 was shown to promote KSHV reactivation from latency (i.e. herpesvirus) by modulating the RLR-dsRNA receptors MDA5 and RIG-1 and preventing IFN production, which restricts viral replication in the absence of ADAR1 [38]. We were initially surprised that in the absence of ADAR1 this signaling pathway does not play a major role in productive HSV-1 infection, but the other dsRNA sensor, PKR, does. However, this is a clear indication that different herepsviruses or viruses at different stages of their replication may rely on the interaction of ADAR1 with different dsRNA sensors. Moreover, studies on HCMV, KSHV, EBV and HSV-1 show that the cells respond differently to infection with different herpesviruses. For example, HCMV infection triggers upregulation of ADAR1 p110 but not p150 [39], on the other hand EBV and KSHV increase both forms [38, 40, 76], while ADAR1 levels do not change in cells infected with HSV-1 (Figure 1) [39]. It would be very interesting to better understand the molecular mechanisms responsible for these differences, but one can speculate that might depend on the dynamics of four main factors: a) immunostimulatory dsRNAs, b) proteome of the infected cell and c) viral modulators of dsRNA sensing. However, there are few comparatives analyzes between different stages of infection or different viruses, so that these differences cannot be adequately addressed at this stage.

Posttranscriptional modifications, including ADAR-mediated A-to-I editing, of viral transcripts may play an important role in RNA recognition by different dsRNA sensors, but based on several lines of evidence, it can be predicted that A-to-I editing of viral transcripts has limited, if any, relevance to productive HSV-1 infection. First, given the limited editing frequency of miR-H2 during productive infection compared to latency, it can be assumed that other viral transcripts undergo only limited editing during productive infection, and it is unlikely that this affects activation of the dsRNA sensor in the context of robust viral transcription. Second, our study shows that impaired HSV-1 replication in ADAR1-deficient cells can be efficiently rescued by depletion of PKR, a pharmacological inhibitor and ADAR1-caliticaly dead protein, demonstrating that editing is not the limiting factor in ADAR1 deficiency. However, importantly, HSV-1 miRNAs are hyperedited in latency strongly suggesting a role of ADAR proteins in the regulation of latent infection [44, 45].

The key questions are why PKR and not MDA5 or other dsRNA sensors are sufficient to rescue viral replication in ADAR1-deficient cells, and which transcripts trigger the activation of PKR in HSV-1-infected cells. The functions of MDA5, RIG-I and PKR are not redundant, and these proteins bind different RNA structures and motifs in a largely sequence-independent manner (Rehwinkel and Gack, 2020), but little is known about the exact endogenous dsRNAs that activates these proteins, and it is possible that they represent different RNA populations. Thus, it is possible that PKR, but not MDA5, preferentially binds viral IE/E transcripts and that activation of MDA5 is downstream of the initial events. Recently, Chiang at al. have shown that in the late phase of infection (16 h.p.i.) host transcripts rather than viral transcripts are enriched in association with RIG-I, namely 5S ribosomal RNA (rRNA) pseudogene transcripts [77]. Furthermore, one of these transcripts, RNA5SP141, has been shown to be relocalized from the nucleus to the cytoplasm during infection and that silencing of this transcript strongly attenuates the antiviral response to HSV-1. In our study, we analyzed PKR-coimmunoprecipitated RNAs at an earlier time point (6 h.p.i.) that coincided with PKR activation and found that, in contrast to the isotope-antibody control, all viral transcripts tested were enriched. Given that vhs released from tegument depletes the vast majority of host transcripts, our results suggest that PKR binds viral transcripts and triggers antiviral signaling. PKR, in contrast to RIG-I, which binds dsRNA end structures, in particular the blunt end and 5’-triphosphate group (5’ppp), is activated by perfect dsRNAs, requiring a minimum of ∼33 bp for activation (Manche et al. 1992)[78], but also by a number of RNAs containing secondary structural imperfections (bulges, internal loops, as well as short double-stranded RNAs with single-stranded tails)[79]. We predicted the structure of a number of viral transcripts (all IE, selected E and L transcripts), and, not surprisingly, all transcripts tested were predicted to contain dsRNA regions or dsRNA irregularities (not shown), suggesting that these transcripts may indeed activate PKR.

Another possibility is that during HSV-1 infection the activation threshold for PKR is lower than for MDA5 in the absence of ADAR1 p150. Furthermore, although our studies in two cell lines (derivatives of HEK293 and A549) demonstrate that PKR signaling was mainly activated and responsible for the reduced replication in the absence of ADAR1, further studies are needed to determine their interdependence and to exclude the possibility that other dsRNA sensors such as ZBP1, RIG-I and LGP2 also contribute to the innate immune response to viral dsRNA.

Another intriguing hypothesis that arises from our study is that HSV-1 evolved in the presence of the modulator of dsRNA sensors, which allowed the virus to delay the expression of its own inhibitors of these pathways, such as ICP34.5, US11, vhs, all of which are encoded by late genes. In support of this, viral mutants expressing US11 (binds dsRNA and directly or/and indirectly prevents the activation of PKR and also other dsRNA sensors OAS, MAD5 and RIG-I [57, 58, 80], under the control of the IE promoter restores protein synthesis and precludes shutoff of protein synthesis by blocking the phosphorylation of eIF2α triggered by virus lacking ICP34.5. It would be very interesting to test whether the provision of US11 would compensate also for the ADAR1 deficiency. Activation of PKR was also observed in cells infected with another viral mutant, vhs-deficient HSV-1 [63, 81], in which higher levels of dsRNA accumulate compared to WT virus, but in contrast to our observation, activation of PKR by ACV was largely prevented. These results suggest that PKR is activated during IE and E times of infection, probably by viral dsRNAs, and that ADAR1 is required to prevent its over-activation. In turn, vhs contributes to this process by depleting dsRNA, and by other mechanisms, but its function is not sufficient to complement ADAR1 deficiency.

We show that in ADAR1-deficient cells, HSV-1 infection triggers activation of PKR, which in turn phosphorylates eIF2α, leading to translational arrest. Although global nascent protein synthesis was reduced compared to uninfected cells, we were surprised that the levels in ADAR1 WT and KO cells were comparable. This result can be explained by the low resolution of the assays used and the relatively small difference in viral replication (up to 10-fold) at the high MOI at which the experiment was performed. Interestingly, we observed that translational arrest affected different viral transcripts differently. Late and early proteins were strongly affected, and this effect could be due to the inhibition of IE proteins. Curiously, we observed that the IE proteins ICP0 and IPC4 were synthesized to a similar extent in the early phase of infection (up to 7 h.p.i.), and in the late phase we observed a stronger decrease of ICP0 but not of ICP4. Important to note, it has been previously observed previously that in vhs selectively impairs translation of viral true late mRNAs, sparing other mRNAs, including ICP4. Clearly additional studies are required to address this; however, it is possible that some HSV-1 transcripts are less sensitive to eIF2α-p translational arrest and selective translation, similar to ATF4.

In addition, we show that HSV-1 infection stimulates direct ADAR1-PKR association, similar to what has been observed after IFN stimulation or infection with RNA viruses. The exact dynamics of these interaction is not well understood; however, it has been shown that requires ADAR1 dsRBD domain and that Z binding domains are dispensable [74, 82]. More research is needed to understand these processes, but our study suggests similar, if not the same, pathways of PKR activation upon IFN stimulation and HSV-1 replication.

In summary, our study identified ADAR1 as a host virulence factor required for efficient HSV-1 infection and a potential target for drug development. Specific inhibitors that target the surfaces that interacts with the dsRNA sensors, rather than the catalytically active site, may allow cells to mount a stronger immune response and limit viral replication in addition to the standard nucleoside analog therapy. Importantly, viruses, that are resistant to standard nucleoside analogs, are also likely to be sensitive to ADAR1 inhibitors and much less likely to develop resistance.

## Material and Methods

### Cells, viruses, plasmids

ADAR1 WT and ADAR1 KO (derivatives of HEK293 cells were generously provided by Jonathan Maelfeit (CRIG, Ghent University) [17, 65]; African green monkey Vero cells (ATCC, CCL-81), epithelial cervical adenocarcinoma cells HeLa (ATCC, CRM-CCL-2), ATCC, lung fibroblast MRC-5 cells (ATCC, CCL-171), epithelial lung carcinoma cells A549 (ATCC, CRM-CCL-185), primary foreskin fibroblast cells (HFF) (kind gift from Stipan Jonjić, Faculty of Medicine, University of Rijeka) were cultured in Dulbaco’s Modified Eagle Medium (PAN Biotech) with 10% fetal bovine serum (PAN Biotech), 100μg/μL Penicillium-Streptomycin (Capricorn), 2mM L-Glutamine (Capricorn) and 1mM Sodium Pyruvate (Capricorn) under standard culture conditions in a humidified incubator at 37C and 5% CO2. Herpes simplex virus 1 (HSV-1) strain KOS (kindly provide by Donald M. Coen Harvard Medical School) and HSV-2 a thymidine kinase (TK)-negative mutant of HSV-2 strain 186syn+,186ΔKpn (HSV-2) (kindly provided by David M. Knipe, Harvard Medical School) [83] were propagated and titrated on Vero cells as previously described [47]. The plasmid DNA pmGFP-ADAR1-p110 (Addgene #117928), pmGFP-ADAR1-p150 (Addgene #117927) contain the ADAR1 gene in pcDNA3.1 expressing p110 and p150 forms, respectively and were constructed by Kumiko Ui-Tei Laboratory [84], pc-FLAG-ICP34.5 contains the ICP34.5 gene (ORF ICP34.5 HSV-1 strain 17) fused to 3x FLAG at N-term was constructed in the laboratory of Donald M. Coen [85], and pCDNA3.1. (ThermoFischer) was prepared using the NucleoBond XTRA Midi kit (Macherey-Nagel # 740410.50) according to the manufacturer’s instructions.

### Infections and plaque assay

The indicated cells were seeded in TC grade plates 24 hours prior to infection. For experiments with HEK293A WT or ADAR1 KO, the plates were coated with 1:200 diluted Matrigel (Corning) for 1 hour before seeding. The next day, the cells were infected with HSV-1 strain KOS at the indicated MOI, and one hour after infection, the infectious medium was replaced with fresh medium. At the indicated time points after infection, the cell-free supernatants were collected and the viral yield was determined using the plaque assay as previously described [86]. Briefly, Vero cells were seeded in 24-well plates at a density of 1.5x10^5^ cells per well and incubated overnight to obtain a monolayer with ∼100% confluence. At the time of titration, the grown medium was removed and the cells were infected with 250uL of a 10-fold serial dilution of the collected supernatant from the infection experiments. After one hour, the infectious medium was removed and the cells were overlayed with methylcellulose media. After 3 days, the cells were fixed with a fixative (5% methanol and 10% acetic acid) for at least 2 hours. The fixative solution was removed and the cells were stained with 5% Giemsa in 1X PBS for at least 2 hours. Excess staining was removed by washing, and the number of plaques was determined. Titers were calculated as plaque-forming units per ml.

### Transfections

The indicated cells were seeded into the TC grade plates 24 hours prior to transfection and transfected with the plasmid DNA using Lipofectamine 2000 (Invitrogen # 11668-027) according to the manufacturer’s instructions. For downregulation of target proteins, cells were seeded in the indicated plates and transfected the next day with 20 pmol siRNAs (supplement list 1) using RNAiMAX (Invitrogen) according to the manufacturer’s instructions. Transfected cells were harvested for analysis or infected with HSV-1 at the indicated MOI 24 hours after transfection. The infection samples for the analysis were taken at the times indicated. For the siRNA screen, cells were seeded in 48-well plates at a density of 10^5^/well and transfected 24 hours later with 20 pmol/well of the indicated siRNAs using RNAiMAX according to the manufacturer’s instructions. 24 hours post-transfection, cells were infected with HSV-1 at MOI 1, and supernatants were collected 24 hours post-infection for titration. The experiment was performed in biological quadruplicates.

### Western blot

For Western blot analysis cells were lysed in RIPA buffer (150 mM NaCl, 1% NP-40, 0.5% Na deoxycholate, 0.1% SDS, 50 mM Tris (pH 8.0)) supplemented with Complete protease inhibitor (Roche) and mixed with 4x Laemelli buffer (Biorad). The proteins were separated in 10% denaturing PAGE and transferred to a nitrocellulose membrane (Machery-Nagel) and Western blotting was performed as previously described [87] using indicated antibodies (Supplement list 2). Proteins were visualized with Amersham ECL (Cytiva) or Supersignal West Femto Detection Reagent (ThermoFischer Sc.) in Chemidoc (BioRAD).

### Nucleic Acid Extraction and RT-qPCR

For RNA analysis, mock-infected and infected cells were harvested at the indicated time points and lysed with TRIreagent (Invitrogen), and total RNA was extracted according to the manufacturer’s protocol. For analysis of cellular and viral transcripts, quantitative reverse transcription-PCR (qRT-PCR) was performed as previously described with modifications. In brief, extracted total RNA was converted to cDNA with High-Capacity kit (Applied Biosystem) according to the manufacturer’s instructions and used to amplify target genes with specific primers (Supplement list 3) and low rox SybrMix (PCR Biosystems) according to the manufacturer’s instructions. All transcripts were normalized to the cellular 18S level (dCT) of the mock infected sample in the respective group, additionally the viral transcripts were normalized to the dCT of the specific transcript at 1hpi of the group.

### dsRNA analysis

For detection of dsRNAs, cells were infected with HSV-1 at the indicated MOI and harvested at the indicated time points, and RNA was extracted using the RNAeasy kit (Qiagen) according to the manufacturer’s instructions. For dot-blot analysis, 1μg of total RNA, 1μg/μl and 0.5 μg/μl of both dsRNA positive control RNA (Jena Bioscience) and negative control RNA (Jena Bioscience) were loaded onto a Nytran Supercharge membrane (Whatman) and UV-crosslinked at 1200mJ/cm2. To normalize the loading of the tested samples, the membranes were stained in 0,1 % solution of methylene blue. For the detection of dsRNA, the membranes were pretreated in a blocking solution (DIG High Prime labelling and detection starter Kit II, Roche) according to the manufacturer’s instructions and incubated overnight at room temperature in a solution of the J2 antibody (1:200, Jena Bioscience). After incubation, the membranes were washed 3 times in TBS-T (0.05% Tween-20 in 1x TBS) and incubated with an HRP-positive secondary antibody (Cell-Signaling). RNAs were visualized with Supersignal West Femto Detection Reagent (ThermoFischer Sc.) in Chemidoc (BioRad) and quantified with ImageJ [88].

### Immunoprecipitation

For protein-protein interaction analysis, cells were mock infected, infected with HSV-1 or treated with 10ng/μl Recombinant IFN-beta (R&D Systems) and lysed in RIPA buffer. Cell debris was precleared by centrifugation at 10000 rpm for 10 minutes at 4°C, and the supernatant was transferred to fresh tubes. 1/5^th^ fraction was set aside for total protein analysis, and remaining was incubated with a primary antibody against PKR (1:50 dilution) (Cell Signalling) or an isotype control (Cell Signalling) gently rolled overnight at 4°C. The next day, 50uL Dynabeads G (Invitrogen # 10004D) were added and the tubes were incubated overnight at 4°C with gentle rolling. The next day, the beads were washed five times with RIPA buffer and boiled directly for 7 minutes at 95°C with 4x Laemelli buffer for Western blot analysis or TRIreagent was added for RNA extraction.

### OPP assay

ADAR1 or ADAR1 KO cells were seeded in 12 well plates and infected with HSV-1 at indicated MOI as described earlier [47] [47, 89]. OPP (Jena Bioscience) was added to the culture medium at a concentration of 50 μM for 1 h before collection and at 12hpi cells were fixed with ice-cold methanol for 2 min at −20 °C [89]. After fixation, cells were washed twice with Tris-buffered saline [TBS: 10 mM Tris (pH 7.5), 150 mM NaCl], permeabilized with TBS with 0.2% Triton X-100 for 7 minutes, and washed twice with TBS. Fixed cells were incubated with staining solution for 30 min ([100 mM Tris (pH 8.5), 0.75 mM CuSO4, 75 mM Na-ascorbate, and 20 μM azide conjugated to Alexa Fluor 594 (Molecular Probes)], washed twice with TBS with 0.5% Triton X-100, and incubated with DAPI for 5 min. Cells were rinsed four times with PBS and once with distilled H2O before inclusion in ProLong Gold antifade reagent (Life Technologies). Images were taken with a Zeiss LSM700 confocal laser scanning microscope (Carl Zeiss) using a 40× Plan Apochromat objective, and results were analyzed with the ZEN imaging software package provided by Zeiss. Mean immunofluorescence signal intensity per cell was obtained by dividing the total fluorescence intensity (n = 8–10 fields with 1,500 cells) by the number of cells.

### Statistical analysis

All analysis was performed with Graphpad Prism 8. For comparison of differences between two groups, unpaired Students’s T test was used. Multiple samples in one group was compared with ordinary one-way annova and multiple samples with multiple groups were compared with two-way annova or mixed model. P-values are quantified against control in respective group and are denoted as p>0.05 as ‘ns’, p≤0.05 as ‘*’, p<0.01 as ‘**’, p<0.001 as ‘***’, and p<0.0001 as ‘****’. Unless otherwise indicated, error bar denotes mean ± SD for all figures.

## Acknowledgements

We thank Edward Mocarski for his critical review of the manuscript and valuable discussion. We thank Mary O’Connell and Liam P. Keegan (CEITEC, Brno) for support and reagents. We thank Donald M. Coen and David M. Knipe (Harvard Medical School) for providing reagents. We thank Sara Patačko and Adrian Perhat for technical assistance.

## Founding

The study was supported by Croatian Science Foundation grant (CSF) IP-2020-02-2287 and DOK-2021-02.9152, and University of Rijeka support grant prirod-sp-23-502930 to I.J.; the European Regional Development Fund (“Strengthening the capacity of the Scientific Center of Excellence CerVirVac for research in viral immunology and vaccinology”; KK.01.1.1.01.0006), CSF IP-2022-10-9967 to SV. UR was supported by grants of the Italian Ministry of University and Research (MIUR): 202292P4R7 and P2022JEEMT. Research in the J.M. group was supported by an Odysseus II Grant (G0H8618N), EOS INFLADIS (40007512), and a junior research grant (G031022N) from the Research Foundation Flanders (FWO).

